# Membrane Topology of UbiA Prenyltransferase Domain-Containing Protein-1 (UBIAD1), a Novel Regulator of Cholesterol Homeostasis

**DOI:** 10.1101/2023.03.02.530834

**Authors:** Brittany Johnson, Dong-Jae Jun, Russell A. DeBose-Boyd

**Affiliations:** Department of Molecular Genetics, University of Texas Southwestern Medical Center at Dallas

**Author notes:** Address correspondence to: Dr. Russell A. DeBose-Boyd, Department of Molecular Genetics, University of Texas Southwestern Medical Center, 5323 Harry Hines Blvd., Dallas, TX 75390-9046.

## Abstract

UbiA prenyltransferase domain containing protein-1 (UBIAD1) is a polytopic membrane-bound enzyme that synthesizes the vitamin K_2_ subtype menaquinone-4 (MK-4) by conjugating the prenyl group of geranylgeranyl pyrophosphate (GGpp) to the aromatic acceptor menadione. The enzyme moonlights as a regulator endoplasmic reticulum (ER)-localized 3-hydroxy-3-methylglutaryl coenzyme A reductase (HMGCR), the rate limiting enzyme in the synthesis of sterol and nonsterol isoprenoids. When cells are replete with isoprenoids, UBIAD1 constitutively cycles between membranes of the medial-trans Golgi and the ER. When ER membranes become depleted of GGpp, UBIAD1 becomes trapped in the organelle where it binds to and inhibits the ER-associated degradation (ERAD) of HMGCR. This inhibition permits continued synthesis of nonsterol isoprenoids, even when sterols are abundant. The resultant accumulation of GGpp in the ER causes dissociation of the HMGCR-UBIAD1 complex, which allows maximal ERAD of HMGCR and translocation of UBIAD1 to the Golgi. These findings disclose a novel GGpp sensing mechanism that allows for metabolically-regulated, intracellular trafficking of UBIAD1. However, the mechanism for this GGpp-induced transport remains to be determined. In the current study, we use cysteine derivatization and protease protection assays to determine the membrane topology of UBIAD1. These findings are key to the determination of mechanisms through which GGpp modulates the intracellular trafficking of UBIAD1 and ultimately, the ERAD of HMGCR.

## Introduction

UbiA prenyltransferase domain-containing protein-1 (UBIAD1) is a member of the UbiA prenyltransferase superfamily of polytopic, integral membrane enzymes found in all forms of life. These enzymes transfer isoprenyl groups to aromatic acceptors, generating molecules such as ubiquinones, vitamin E, and vitamin K.^1^ In mammalian cells, UBIAD1 synthesizes the vitamin K_2_ subtype menaquinone-4 (MK-4) by using the nonsterol isoprenoid geranylgeranyl pyrophosphate (GGpp) to prenylate 1,4-naphthoquinone menadione (MD or vitamin K_3_) derived dietary phylloquinone (PK or vitamin K_1_)^2^,^3^. UBIAD1 is multifunctional as indicated by the association of missense mutations in the human *UBIAD1* gene with Schnyder corneal dystrophy (SCD). The rare, autosomal-dominant disease is characterized by the bilateral opacification of the cornea, which results in part, from the overaccumulation of cholesterol.^4^,^5^,^6^,^7^,^8^ Our group showed that SCD-associated UBIAD1 blocks the sterol-accelerated, endoplasmic reticulum (ER)-associated degradation (ERAD) of 3-hydroxy-3-methylglutaryl coenzyme A (HMG CoA) reductase (HMGCR), which catalyzes the reduction of HMG CoA to mevalonate.^9^ This reaction has been long-recognized as the rate-limiting reaction in the branched pathway that produces cholesterol and nonsterol isoprenoids such as GGpp.^10^,^11^,^12^,^13^

Sterol-accelerated ERAD is one of several mechanisms that converge on HMGCR to ensure that cells continuously produce GGpp and other essential nonsterol isoprenoids but avoid the toxic overaccumulation of cholesterol.^14^ This ERAD is initiated by the build-up sterols in membranes of the ER, which causes HMGCR to bind to ER membrane proteins called Insig-1 and Insig-2.^14^ At least three Insig-associated E3 ubiquitin ligases mediate ubiquitination of HMGCR, marking it for extraction across the ER membrane by the ATPase valosin-containing protein (VCP)/p97 and its ubiquitin-binding cofactors.^15^,^16^,^17^ Extracted HMGCR is then dislodged from ER membranes into the cytosol and delivered to 26S proteasomes for proteolytic degradation.^18^,^19^,^20^ Maximal ERAD of HMGCR requires the addition to cells of geranylgeraniol (GGOH), the alcohol derivative of GGpp.^20^ Current evidence indicate that one or more kinases convert GGOH to GGpp, which in turn, augments membrane extraction of ubiquitinated HMGCR^42^.

Sterols also cause a fraction of HMGCR molecules to bind UBIAD1. ^21^ This binding inhibits a post-ubiquitination step in HMGCR ERAD, allowing production of small amounts of mevalonate that are preferentially incorporated into GGpp and other nonsterol isoprenoids.^22^ When GGpp builds up in membranes of the ER, the isoprene binds to UBIAD1 and triggers its dissociation from HMGCR. Dissociation of the HMGCR-UBIAD1 permits the maximal ERAD of HMGCR.^23^,^24^,^25^ The structural analyses of archaeal UbiA prenyltransferases revealed that residues corresponding to SCD-associated mutations in human UBIAD1 cluster around the active site, indicating a reduced affinity for GGpp.^26^,^27^ Indeed, SCD-associated UBIAD1 is resistant to GGpp-induced dissociation from HMGCR and inhibits ERAD in a dominant-negative manner.^9^

During our studies, we discovered that GGpp also stimulates translocation of UBIAD1 from the ER to the medial-trans Golgi.^9^ This finding formed that basis for a model that predicts UBIAD1 constitutively cycles between membranes of the ER and Golgi when cells are replete with GGpp. When UBIAD1 senses a decline in GGpp embedded in ER membranes, the enzyme becomes trapped in the organelle where it binds to and inhibits ERAD of HMGCR to enhance synthesis of GGpp and other nonsterol isoprenoids. This novel sensing mechanism is defective in SCD, which is indicated by the finding that SCD-associated UBIAD1 remains sequestered in the ER of cells replete with GGpp.^9^ The resultant inhibition of HMGCR ERAD causes enhanced synthesis and intracellular accumulation of cholesterol in cultured cells and tissues of mice expressing SCD-associated UBIAD1.^28^

Anterograde transport of proteins from ER to Golgi occurs through their incorporation into transport vesicles that bud from ER membranes. These vesicles are generated by coat protein complex (COP)-II, which is composed of five cytosolic proteins: the GTPase Sar1, Sec23, Sec24, Sec13, and Sec31. The recruitment of GTP-bound Sar1 to ER membranes initiates formation of COPII vesicles. Membrane-bound Sar1 recruits Sec23 and Sec24 heterodimers. Sec23 directly associates with Sar1, whereas Sec24 interacts with selects cargo proteins. Sar1 and Sec23/24 subsequently recruit the heterotetrametric Sec13/31 complex that forms the outer COPII coat and causes budding of vesicles from the ER. Budded vesicles soon shed their COPII coat and fuse with cis elements of the Golgi.^29^ Methods that reconstitute *in vitro* formation of COPII vesicles have been established. Using these methods, we found that GGpp stimulated incorporation of UBIAD1 into ER-derived transport vesicles.^9^ Importantly, SCD-associated UBIAD1 fails to become incorporated into these transport vesicles in the presence of GGpp. These findings have led us to propose that the binding of GGpp to UBIAD1 induces a conformational change in the protein that exposes a binding site for COPII, which permits its incorporation into transport vesicles. In order to determine how UBIAD1 associates with COPII, we need to identify the COPII binding site of UBIAD1. To facilitate the identification of this COPII binding site, it is essential to delineate the topology of UBIAD1 in membranes. Here, we use the combination of protease protection and cysteine derivatization to show that UBIAD1 is integrated into membranes by 9 membrane-spanning helices separated by short cytosolic and luminal loops.

## Results

We analyzed the topology of human UBIAD1 using six different algorithms (**Figure 1A**, top)^30^. Four of the six algorithms predicted that UBIAD1 contains nine transmembrane domains, one algorithm predicted seven transmembrane domains, whereas the other predicted six transmembrane domains with a signal peptide at the N terminus. The hydropathy plot of the amino acid sequence of human UBIAD1 generated by the TOPCONS algorithm predicted UBIAD1 contains nine transmembrane domains that are depicted by peak regions of hydrophobicity (**Figure 1A**, bottom). Based on data from the experiments described here, we suggest a membrane topology for UBIAD1 depicted in **Figure 1B**. According to our model, the N-terminus of UBIAD1 (corresponding to amino acids 1-45) is cytosolic, while the C-terminus of the protein extends into the lumen. The N- and C-termini of UBIAD1 are separated by nine membrane-spanning helices. Our predicted model for the topology of UBIAD1 is not consistent with the hydropathy plot of the protein shown in **Figure 1A**.

**Figure 1.**
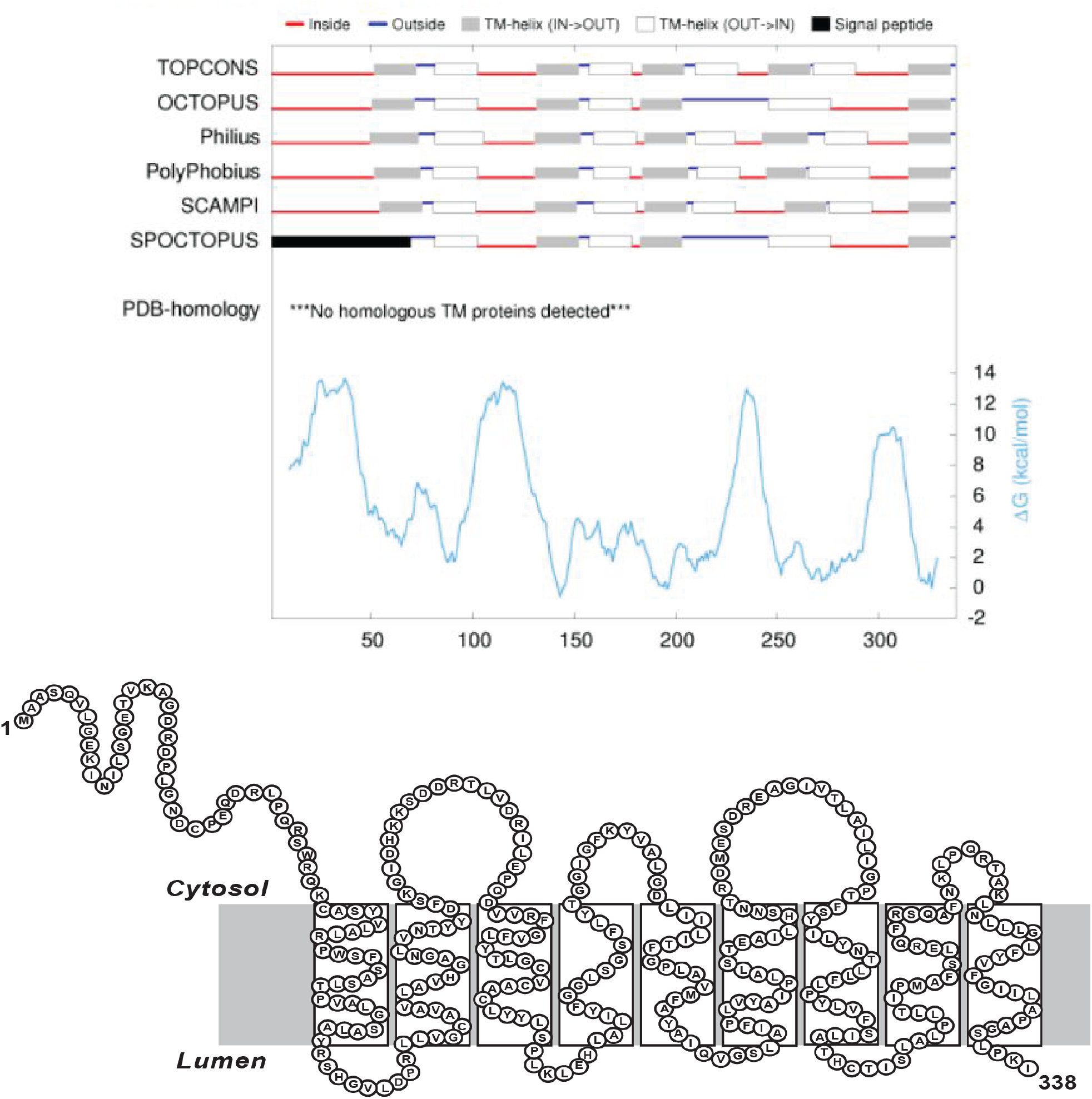
Hydropathy plot and predicted membrane topology of human UBIAD1. **A**, Hydropathy plot of UBIAD1:Seven different predicted topologies and the algorithm used to predict them are displayed at the top of the figure, the residue-specific Gibbs Free Energy change was calculated using the web server TOPCONS. The putative transmembrane helices are indicated in grey and white, the cytosolic regions are indicated in blue, the luminal regions are indicated in red, and the signal peptide are indicated in black. On the bottom of the graph is the Gibbs Free Energy change plotted against the amino acid position. **B**, Topology model for UBIAD1 based on the data in this manuscript.

In **Figure 2**, we began to evaluate our predicted model for the membrane topology of UBIAD1 by preparing expression plasmids encoding human UBIAD1 containing an N-terminal Myc tag (pCMV-Mycx3-hUBIAD1) or a C-terminal FLAG tag (pCMV-UBIAD1-FLAG3x). We next transfected transformed human fibroblasts (designated SV-589, ref) with either pCMV-Myc-UBIAD1 (**Figure 2A**) or pCMV-UBIAD1-FLAG (**Figure 2B**). The cells were subsequently harvested for preparation of lysates from which we isolated intact membrane fractions. The membrane fractions were then incubated with increasing concentrations of trypsin in the absence or presence of the detergent NP-40. The proteolytic reactions were terminated and samples were fractionated by SDS-PAGE and subjected to immunoblot analysis with anti-Myc and anti-FLAG antibodies. In the absence of trypsin, a single 28 kDa band corresponding to Myc-UBIAD1 was detected (**Figure 2A, lane 1**); trypsinolysis caused the complete disappearance of this band (**Figure 2A, lanes 2-4**). Similar results were obtained in the presence of NP-40 (**Figure 2A, lanes 5-8**). A similar 28 kDa band was observed for UBIAD1-FLAG (**Figure 2B, lane 1**). However, trypsin did not cause complete disappearance of the band but rather led to the generation of a 13 kDa fragment (**Figure 2B, lanes 3 and 4**) that was digested in the presence of NP-40 (**Figure 2B, lanes 7 and 8**). To analyze membrane integrity, we probed for the ER-resident protein calnexin using an antibody directed against its large lumenal domain (**Figures 2A** and **2B**). The results show that the lumenal domain of calnexin resisted trypsin digestion in the absence of NP-40 (**Figures 2A and B, lanes 1-4**), but not in the presence of the detergent (**Figures 2A** and **B, lanes 5-8**). Thus, membrane vesicles generated in our studies were sealed and impermeant to trypsin in the absence of NP-40. Considered together, the results of **Figure 2** indicates that the N-terminus of UBIAD1 extends into the cytosol, whereas the C-terminal end of the protein localizes to the lumen.

**Figure 2.**
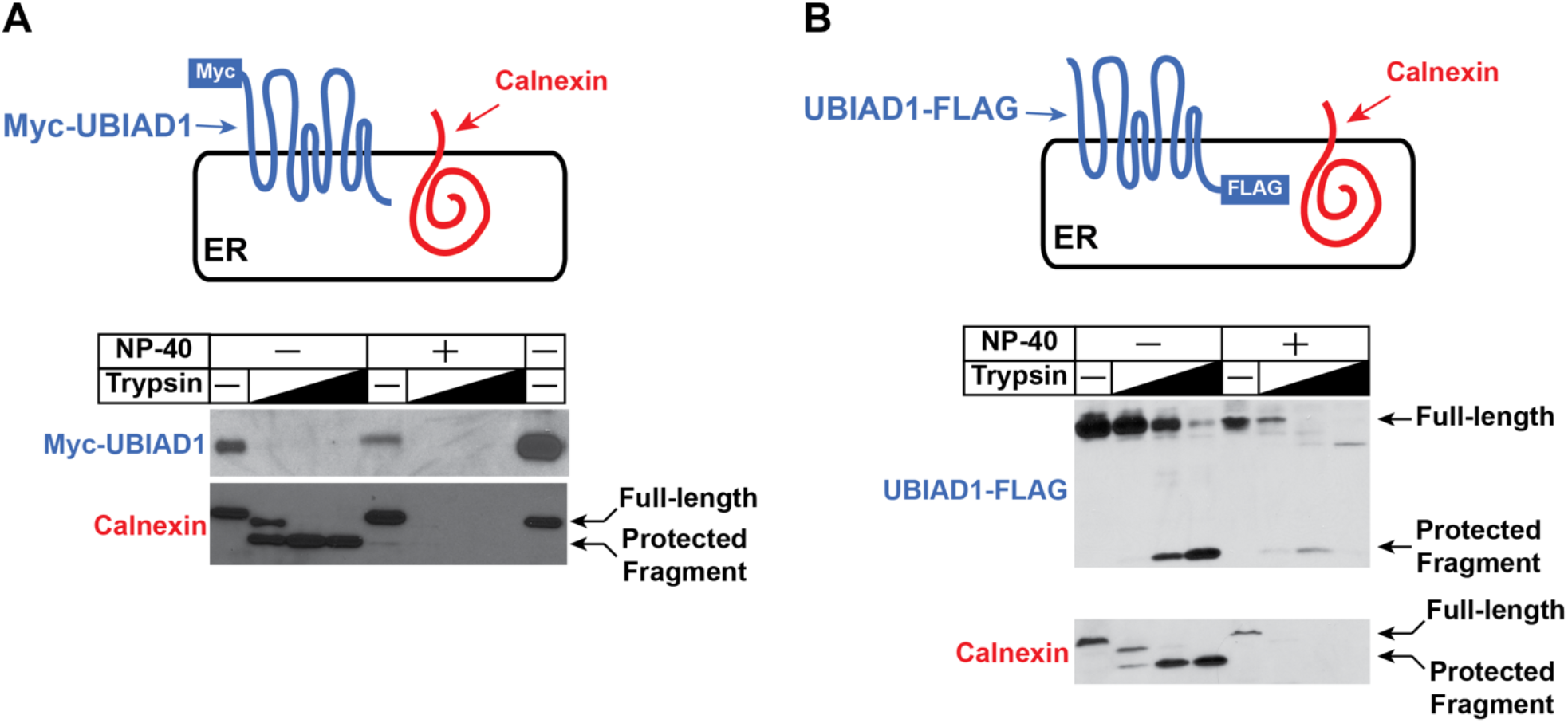
Membrane orientation of the NH2 terminal and the COOH-terminal of UBIAD1 as determined by trypsin proteolysis. **A**, A schematic illustration of the fusion protein expressed from cDNA used in this figure. The expression plasmid pCMV-Myc-UBIAD1, which encodes human UBIAD1 containing a single copy of a Myc epitope at the N-terminus under transcriptional control of the cytomegalovirus (CMV) promoter, was previously described. The expression plasmid pCMV-UBIAD1-FLAG, which encodes human UBIAD1 containing a single copy of a FLAG epitope at the C-terminus under transcriptional control of the cytomegalovirus (CMV) promoter, was also previously described. SV589 cells were transfected with 5 μg of pCMV-MYC-UBIAD1 or pCMV-UBIAD1-FLAG in 100 mm dishes. Microsomes were harvested without protease inhibitors. An equal concentration of microsomal protein was suspended in a total volume of 56 μl in Buffer A and was incubated with the indicated amounts of trypsin for 30 min at 30 °C in the absence or presence of the detergent NP-40. Reactions were terminated by the addition of loading buffer and heat inactivation at 95 °C for 10 min.

In **Figure 3**, we implemented a method of cysteine derivatization called PEGylation to further examine the membrane topology of UBIAD1.^31^ UBIAD1 contains seven cysteine residues. To facilitate the cysteine derivatization experiments, we first prepared an expression plasmid encoding a mutant version of Myc-UBIAD1 in which all seven cysteines were mutated to serine residues; cysteine-less UBIAD1 is designated Myc-UBIAD1 (Cys-Null). Next, we introduced into Myc-UBIAD1 (Cys-Null) single cysteine residues at thirty-one positions, as shown in **Figure 3**. For cysteine derivitization experiments, we transfected these plasmids into CHO-K1 cells. Sealed membrane vesicles were isolated and pretreated with a membrane impermeable cysteine derivitization agent Mal-Peg. The sealed membranes were then treated with detergent before being subjected to SDS-PAGE and blotted with an antibody against MYC. In the absence of treatment, a single 28 kDa band was observed (**Figure 3, lane 1**). When sealed membranes were harvested from cells that were transfected with the Cys-Null mutant there was no detectable PEGylation despite treatment with mal-peg and detergent therefore a single 28 kDa band was observed (**Figure 3, lanes 2-5**). When sealed membranes were harvested from cells that were transfected with plasmid encoding wild type MYC-UBIAD1, multiple bands were detected due to the seven cysteine residues being pegylated. The band shift became more pronounced with the addition of detergent (**Figure 3, lanes 6-9**). When sealed membranes were harvested from cells that were transfected with a single cysteine mutant of UBIAD1, there was one of three possible results 1) cytosolic cysteines would be PEGylated in the absence of detergent revealing two bands at 28 kDa and 35 kDa 2) luminal cysteines would be PEGylated in the presence of a weak detergent revealing two bands at 28 kDa and 35 kDa or 3) sterically inhibited cysteines would only become PEGylated in the presence of a strong detergent revealing two bands at 28 kDa and 35 kDa (**Figure 3, lanes 10-13**). Cysteines at residues 31,41,46,75,108,116,125,239,304,305,306,307,308,309,310,311,312 were exposed. Cysteines at residues 57, 61, 87, 126,176,177,179,184,209,273 and 279 were buried. Cysteine at residues 69,145 and 224 were sterically hindered. These results led us to conclude that UBAD1 has nine transmembrane domains.

**Figure 3.**
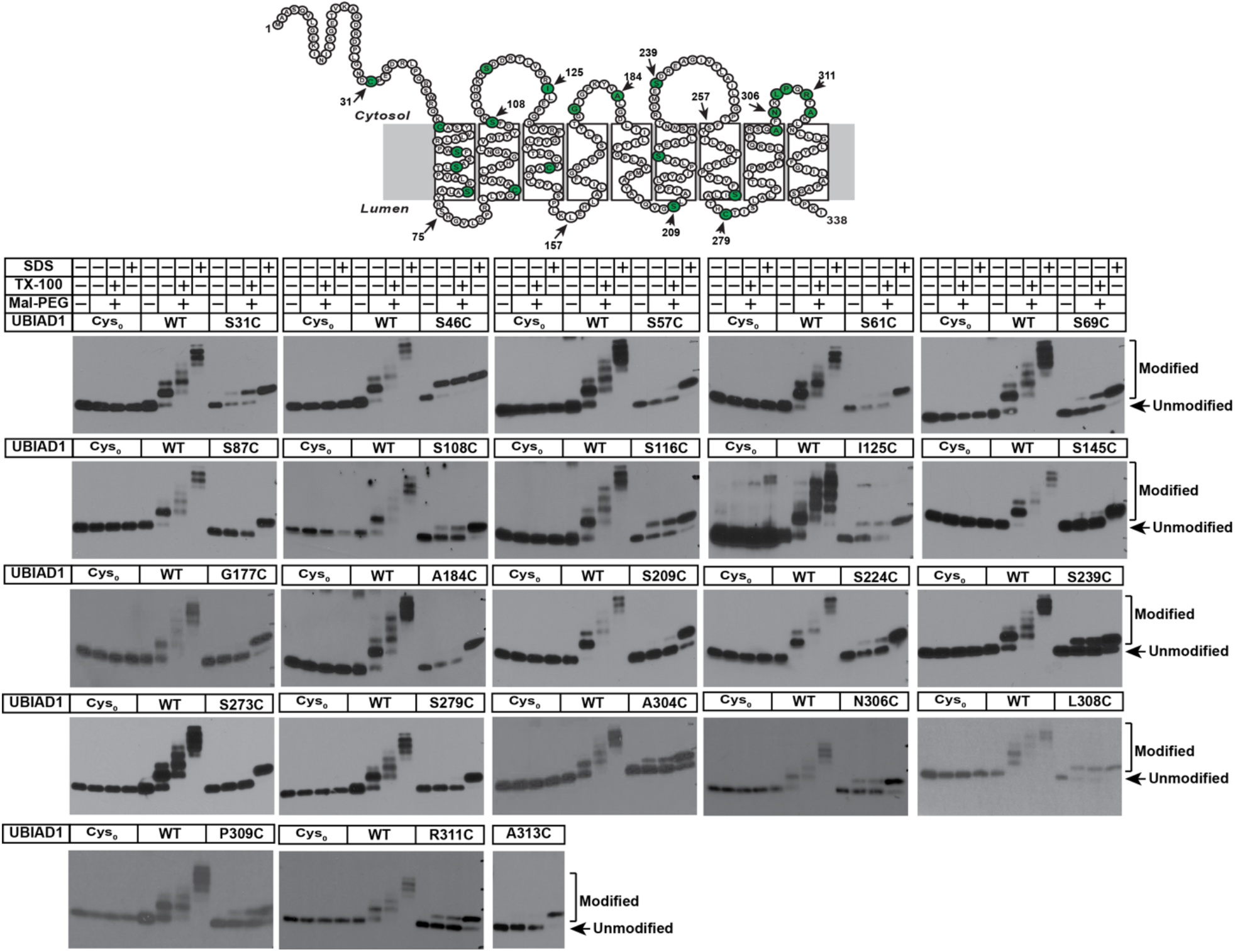
Membrane localization of cysteine residues in UBIAD1 as determined by MAL-dPEG^®^4-(m-dPEG®12)3 derivatization. The PEGylation method was adapted from Lowe et al. CHO-K1 cells were plated at 250K in 10 cm dishes and transiently transfected on Day 2 with 5 μg of plasmid DNA in media containing 5% FCS. Microsomal protein (20 μg) was suspended in a total volume of 54 μl of Buffer A containing protease inhibitors. Microsomes were treated with or without 1% (v/v) Triton X-100 or SDS as indicated. Microsomes were then treated with or without mPEG (catalog number 10406, Quanta Biodesign) (5 mm, 30 min) at 37 °C. Reactions were quenched with dithiothreitol (DTT) (10 mm, 10 min, room temperature). Samples were subjected to Novex 4-20% Tris-Glycine followed by Western blotting.

**Figure 4** examines the subcellular localization of topology mutants in SV-589 (ΔUBIAD1) cells. Consistent with our previous results, most of the topology mutants colocalized with the Golgi protein GM-130 (Pearson correlation coefficient = 0.668) when cells were cultured in isoprenoid-replete medium containing FCS (**Fig. 4, panels 1–36; supplemental Fig. S1**).^9^ The subcellular localization of eleven topology mutants in SV-589 (ΔUBIAD1) cells exhibited a reticular pattern, indicative of localization to ER membranes (**Fig. 4, panels 8,15,17,18,19,28,31,32,34,35,36; supplemental Fig. S1**). Although several topology mutants do not localize in a metabolically regulated fashion, we determined that they are still suitable for mal-peg studies because they mimic the Schnyder Corneal Dystrophy (SCD) phenotype meaning that they maintain normal enzymatic activity and the ability to stabilize HMGCR.

**Figure 4.**
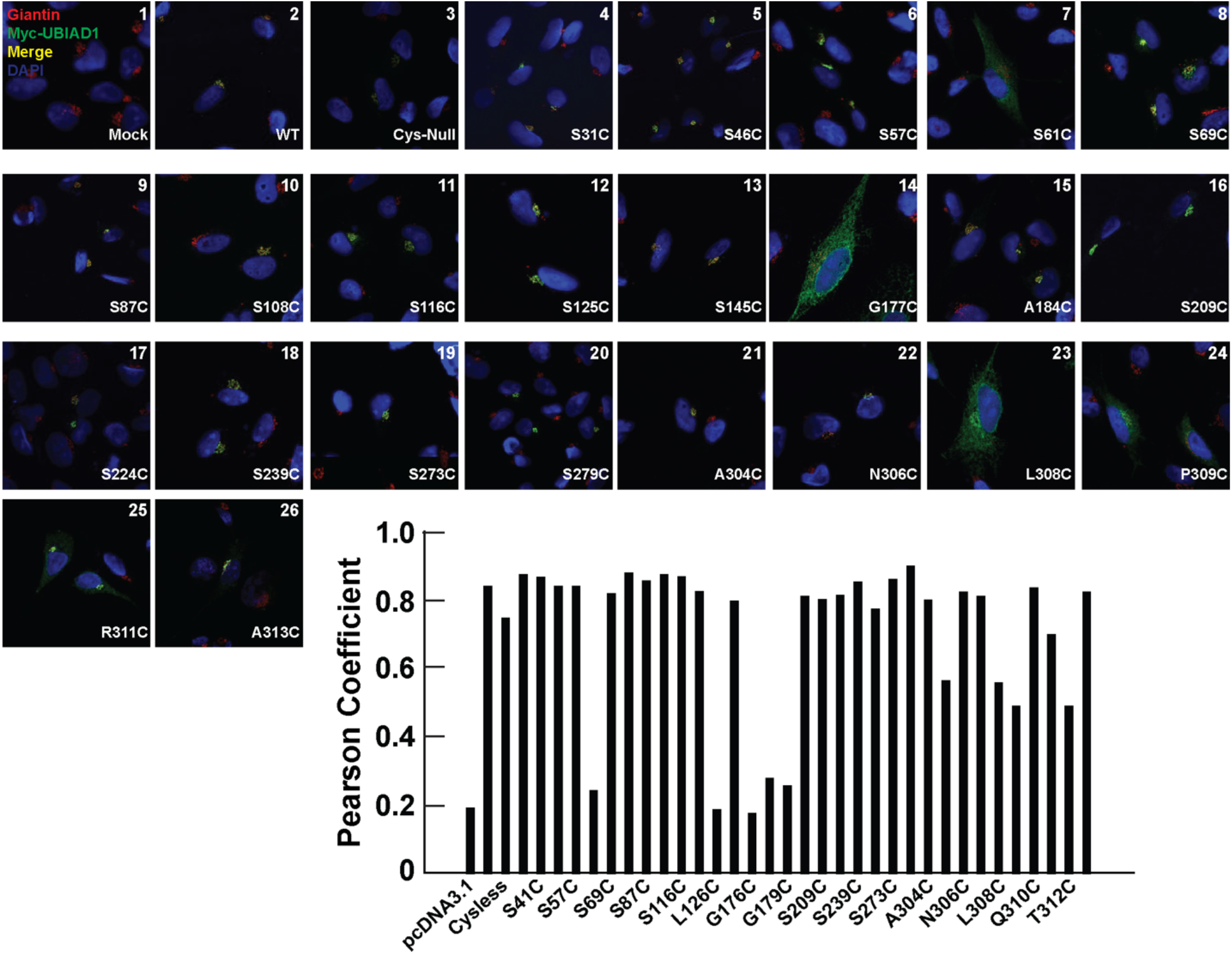
Subcellular localization of UBIAD1 variants used in membrane topology experiments. SV-589 cells were set up for experiments on day 0 at 150K in 60 mm dishes in 5% FCS. On day 1, the cells were transfected with 0.5 μg/ml pf plasmid DNA in 5% FCS. On Day 3, the cells were washed with PBS and subsequently fixed and permeabilized for 15 min in methanol at −20°C. Upon blocking with 1 mg/ml BSA in PBS, coverslips were incubated for 1 hour at room temperature with primary antibodies (rabbit polyclonal anti-GM130 IgG and IgG-9E10, a mouse monoclonal antibody against c-Myc purified from the culture medium of hybridoma clone 9E10 [American Type Culture Collection, Manassas, VA]) diluted in PBS containing 1 mg/ml BSA. Bound antibodies were visualized with goat anti-mouse IgG conjugated to Alexa Fluor 488 and goat anti-rabbit Alexa Fluor 594 (Life Technologies, Grand Island, NY) as described in the figure legends. In addition, coverslips were stained for 5 min with 1 μg/ml 300 nM 4′,6-diamidino-2-phenylindole (Life Technologies) to visualize nuclei. The coverslips were then mounted in Mowiol 4-88 solution (Calbiochem/EMD Millipore, Billerica, MA) and fluorescence analysis was performed using a Plan-Apochromat 63×/1.4 objective (Zeiss, Peabody, MA), an Axio Observer microscope (Zeiss), an Axiocam (Zeiss) color digital camera in black and white mode, and ImageJ software (National Institutes of Health, Bethesda, MD). Brightness levels were adjusted across entire images using ImageJ. The values are the average of 6-10 images (± standard error).

In **Figure 5**, we tested the ability of these topology mutants to stabilize HMGCR in sterol replete conditions. Intact membranes were harvested from SV-589 cells that were co transfected with HMGCR and various topology mutants and then cultured in isoprenoid-replete medium containing FCS overnight. In the absence of UBIAD1 and HMGCR, no band is detected (**Figure 3B, Lane 1**). In the presence of just the HMGCR plasmid, there is a faint band detected at the 37 kDa (**Figure 5, Lane 2**). In the presence of both HMGCR and UBIAD1 (30 kDa), robust expression of HMGCR is observed (**Figure 5, Lane 3**). All the topology mutants stabilize HMGCR in sterol replete conditions except for 249, 257, 308 and 309 (**Figure 5, Lanes 4-13**). Topology mutants 249 and 257 do not stabilize HMGCR in non-sterol isoprenoid replete conditions and topology mutants 308 and 309 hyperstabilze HMGCR in comparison to WT. Residue 249 was not expressed in IF or Mal-Peg studies (data not shown) so we concluded that this mutant is poorly expressed via transient transfection. Residues 257, 305, 311, and 312 were mutated to alanine in a wildtype background and were all found to have normal localization and retained the ability to stabilize HMGCR in isoprenoid replete media (**Supplemental Figure 3, data not shown**). This indicates that the abnormal results of these corresponding topology mutants are possibly due to them being in a cysteineless background. Residue 126 is expressed in Mal-Peg experiments but its localization is not metabolically regulated and it does not synthesize MK-4. This indicates that mutating residue 126 in the cysteineless background generates a catalytically dead enzyme.

**Figure 5.**
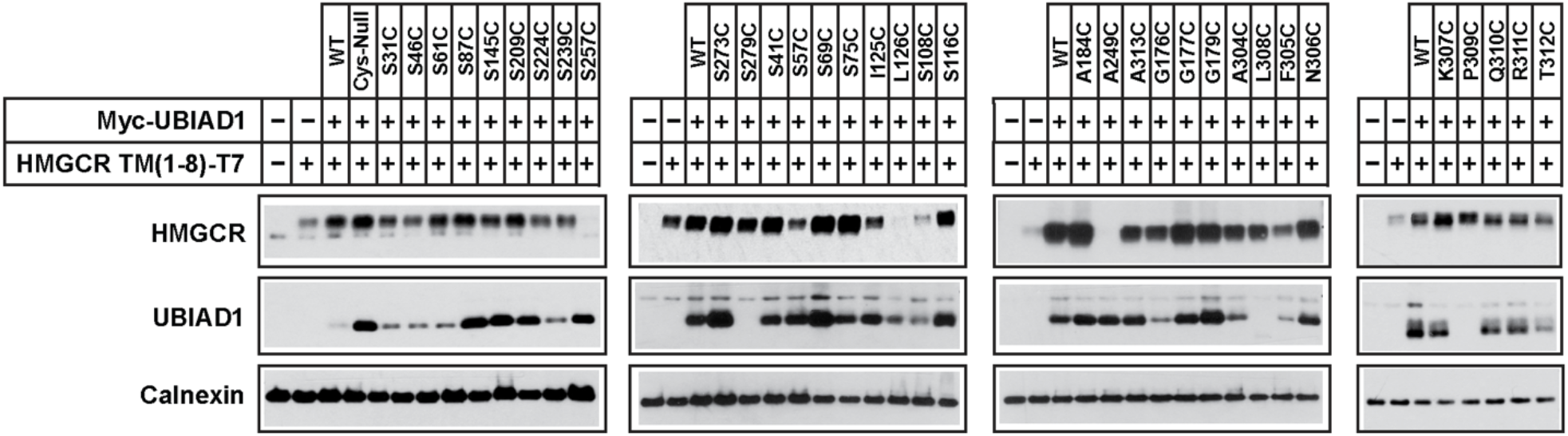
Stabilization of HMG CoA reductase by UBIAD1 variants used in membrane topology experiments. SV-589 cells were set up for experiments at 400K cells per 100-mm dish in media containing 5% FCS. On day 1, cells were transfected with 3 μg/dish of pCMV-HMG-Red(TM1-8)-T7 together with 100 ng of pCMV-Myc-UBIAD1 in media containing 5% FCS. Following incubation for 16 hr at 37°C, cells were harvested and subjected to subcellular fractionation. Aliquots of resulting membrane fractions were then subjected to SDS-PAGE and immunoblot analysis was carried out with anti-T7 IgG (against reductase), IgG-9E10 (against UBIAD1), and anti-calnexin IgG.

In the experiments shown in **Figure 6**, we tested the ability of these topology mutants to synthesize MK-4. Membranes isolated from SV-589 cells that were transfected with a topology mutant were resuspended in buffer and incubated with [3H] menadione. After an overnight incubation, reactions were terminated; lipids were extracted and fractionated by TLC, followed by scintillation counting. A [3H]-labeled product that comigrated with MK-4 on TLC plates was produced by all the reactions that contained membranes transfected with topology mutants except for positions 126 and 249.

**Figure 6.**
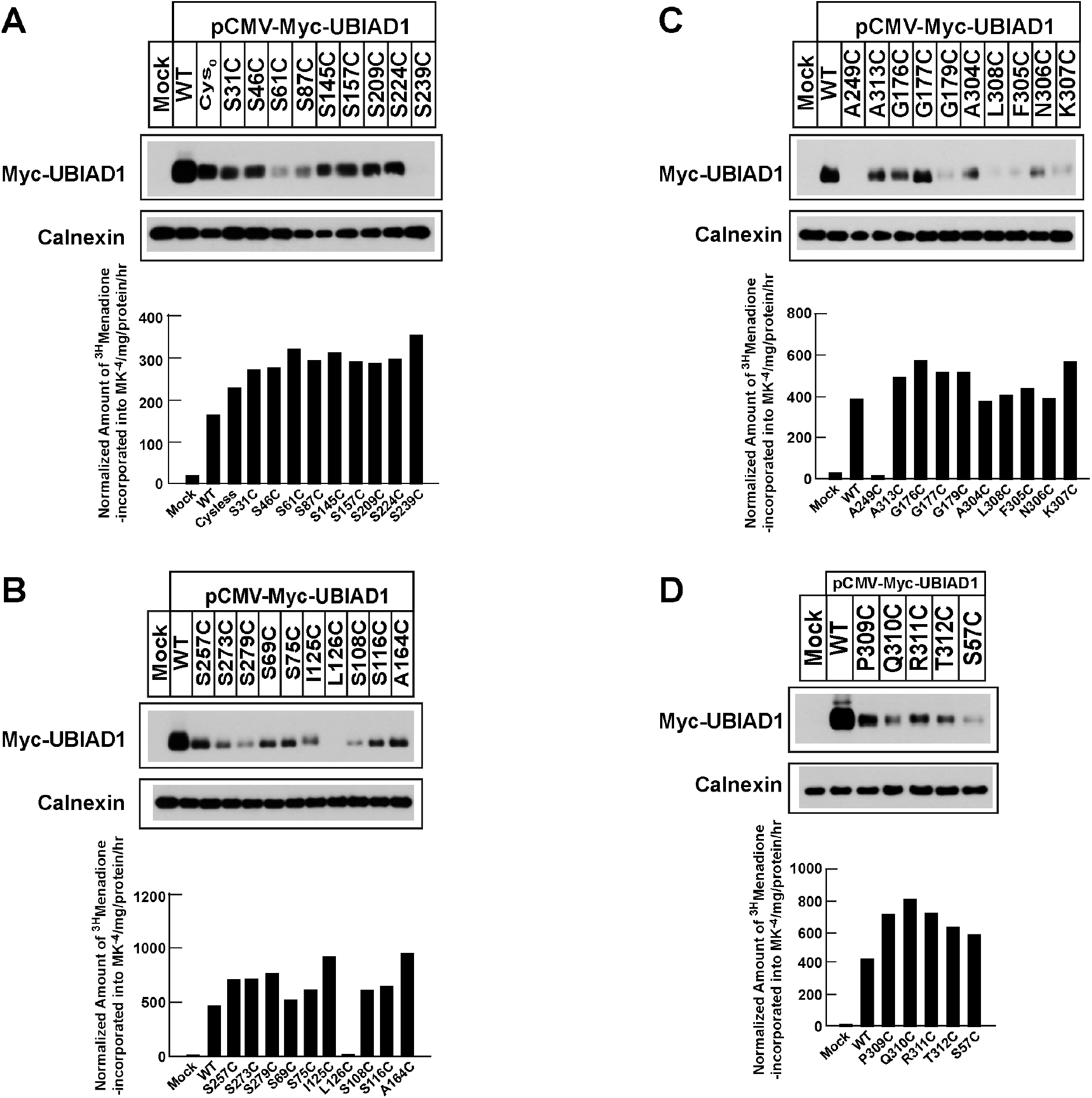
MK-4 synthetic activity of UBIAD1 variants used in membrane topology studies. A-D, To measure [^3^H]MK-4 synthesis in intact cells, we set up cells on day 0 at 250K cells in 60 mm dishes. On day 2, the cells were refed medium A with 5% FCS and 2.5 μCi/ml [3H]menadione (unlabeled menadione was added to achieve a final concentration of 50 nM). Following incubation for 16 h at 37°C, cells were washed twice with PBS containing 2% BSA, followed by an additional wash with PBS. The cells were lysed with 0.1 N NaOH; an aliquot of each lysate was removed for protein determination. The remaining lysates were mixed with recovery solution containing 16 μg/ml MK-4, 0.025 μCi/ml [14C]cholesterol, and 16 μg/ml unlabeled cholesterol and extracted with dichloromethane:methanol (2:1). The resulting lipids were subjected to TLC and incorporation of [^3^H]menadione into MK-4 was determined as previously described.

While conducting IF studies of the topology mutants; we noticed that many of the mutants that did not localize in a metabolically regulated fashion were located at cytosolic loop eight. All these mutants have normal enzymatic activity and stabilize HMGCR in non-sterol isoprenoid replete conditions. This led us to determine that this is a possible cop II binding site. To test this hypothesis, we prepared a series of nine expression vectors encoding human UBIAD1 where each residue on loop eight was mutated to an alanine with a MYC tag added to the N-terminal domain.

In **Figure 7**, we tested the functionality of each topology mutants ability to localize between the ER and the Golgi in a metabolically regulated manner **(Figure 7A)**, stabilize HMGCR in non-sterol isoprenoid replete conditions **(Figure 7B)**, and synthesize MK-4 **(Figure 7C)**. **Figure 7A** examines the subcellular localization of alanine scan mutants in SV-589 (ΔUBIAD1) cells. Consistent with our previous results, most of the alanine scan mutants colocalized with the Golgi protein GM-130 (Pearson correlation coefficient = 0.668) when cells were cultured in isoprenoid-replete medium containing FCS **(Fig. 3A, panels 1–36; supplemental Fig. S1**). The subcellular localization of one alanine scan mutant in SV-589 (ΔUBIAD1) cells exhibited a reticular pattern (P309A), indicative of localization to ER membranes **(Fig. 7A, panel 7**). We tested the functionality of each alanine scan mutant for its ability to synthesize MK-4 **(Figure 7B)** and to stabilize HMGCR in non-sterol isoprenoid replete conditions **(Figure 7C)** as previously described. All of the alanine scan mutants stabilize HMGCR in non-sterol isoprenoid replete conditions and have some level of enzymatic activity.

**Figure 7.**
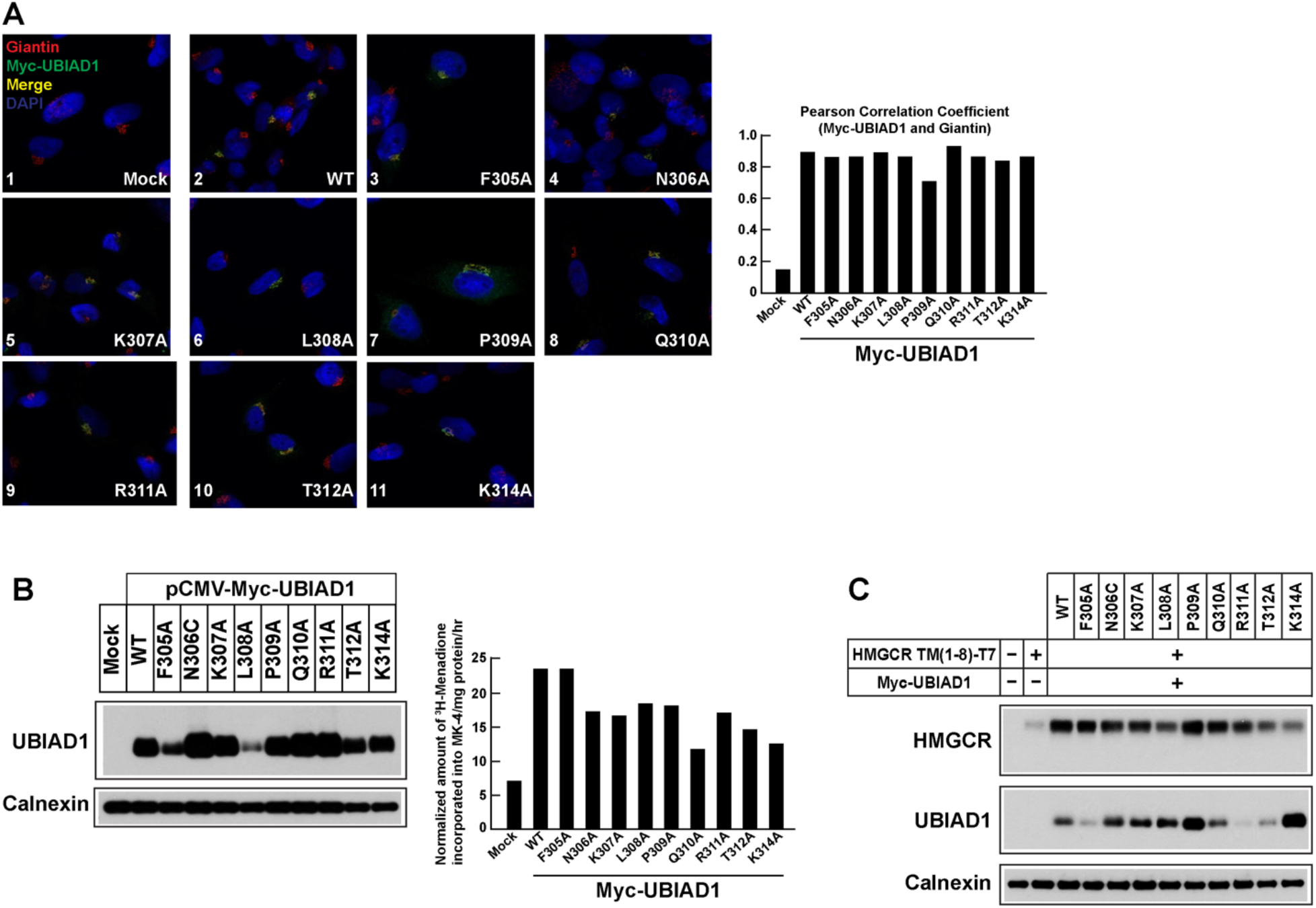
Subcellular localization (A), MK-4 synthetic activity (B), and HMG CoA reductase stabilization (C) of UBIAD1 variants harboring alanine substitutions in cytosolic loop between transmembrane domains 8 and 9. **A**, SV-589 cells were set up for experiments on day 0 at 150K in 60 mm dishes in 5% FCS. On day 1, the cells were transfected with 0.5 μg/ml pf plasmid DNA in 5% FCS. On Day 3, the cells were washed with PBS and subsequently fixed and permeabilized for 15 min in methanol at −20°C. Upon blocking with 1 mg/ml BSA in PBS, coverslips were incubated for 1 hour at room temperature with primary antibodies (rabbit polyclonal anti-GM130 IgG and IgG-9E10, a mouse monoclonal antibody against c-Myc purified from the culture medium of hybridoma clone 9E10 [American Type Culture Collection, Manassas, VA]) diluted in PBS containing 1 mg/ml BSA. Bound antibodies were visualized with goat anti-mouse IgG conjugated to Alexa Fluor 488 and goat anti-rabbit Alexa Fluor 594 (Life Technologies, Grand Island, NY) as described in the figure legends. In addition, coverslips were stained for 5 min with 1 μg/ml 300 nM 4′,6-diamidino-2-phenylindole (Life Technologies) to visualize nuclei. The coverslips were then mounted in Mowiol 4-88 solution (Calbiochem/EMD Millipore, Billerica, MA) and fluorescence analysis was performed using a Plan-Apochromat 63×/1.4 objective (Zeiss, Peabody, MA), an Axio Observer microscope (Zeiss), an Axiocam (Zeiss) color digital camera in black and white mode, and ImageJ software (National Institutes of Health, Bethesda, MD). Brightness levels were adjusted across entire images using ImageJ. The values are the average of 6-10 images (± standard error). **B**, To measure [3H]MK-4 synthesis in intact cells, we set up cells on day 0 at 250K cells in 60 mm dishes. On day 2, the cells were refed medium A with 5% FCS and 2.5 μCi/ml [3H]menadione (unlabeled menadione was added to achieve a final concentration of 50 nM). Following incubation for 16 h at 37°C, cells were washed twice with PBS containing 2% BSA, followed by an additional wash with PBS. The cells were lysed with 0.1 N NaOH; an aliquot of each lysate was removed for protein determination. The remaining lysates were mixed with recovery solution containing 16 μg/ml MK-4, 0.025 μCi/ml [14C]cholesterol, and 16 μg/ml unlabeled cholesterol and extracted with dichloromethane:methanol (2:1). The resulting lipids were subjected to TLC and incorporation of [3H]menadione into MK-4 was determined as previously described. **C**, SV-589 cells were set up for experiments on day 1 400K cells per 100-mm dish in containing 5% FCS. On day 1, cells were transfected with 3 μg/dish of pCMV-HMG-Red(TM1-8)-T7 together with 100 ng of pCMV-Myc-UBIAD1 in media containing 5% FCS. Following incubation for 16 hr at 37°C, cells were harvested and subjected to subcellular fractionation. Aliquots of resulting membrane fractions were then subjected to SDS-PAGE and immunoblot analysis was carried out with anti-T7 IgG (against reductase), IgG-9E10 (against UBIAD1), and anti-calnexin IgG.

To determine if P309A localizes to the Golgi at a slower rate than the rest of the mutants or if P309A is unable to leave the ER, we conducted a time course experiment **(Figure 8)**. In this experiment, we sterol depleted SV589 cells overnight and attempted to restore the Golgi localization of these mutants by treating with geranylgeraniol (GGOH), which becomes phosphorylated to GGpp, over five different time points. We found that all the mutants localize to Golgi within one hour of GGOH treatment **(Figure 8, panels 1-35,37-60**) except for P309A **(Figure 8, panels 31-36**) which took between four to eight hours to localize to the Golgi. This led us to believe that P309A is an important residue for regulating the transport of UBIAD1 between the ER and the Golgi.

**Figure 8.**
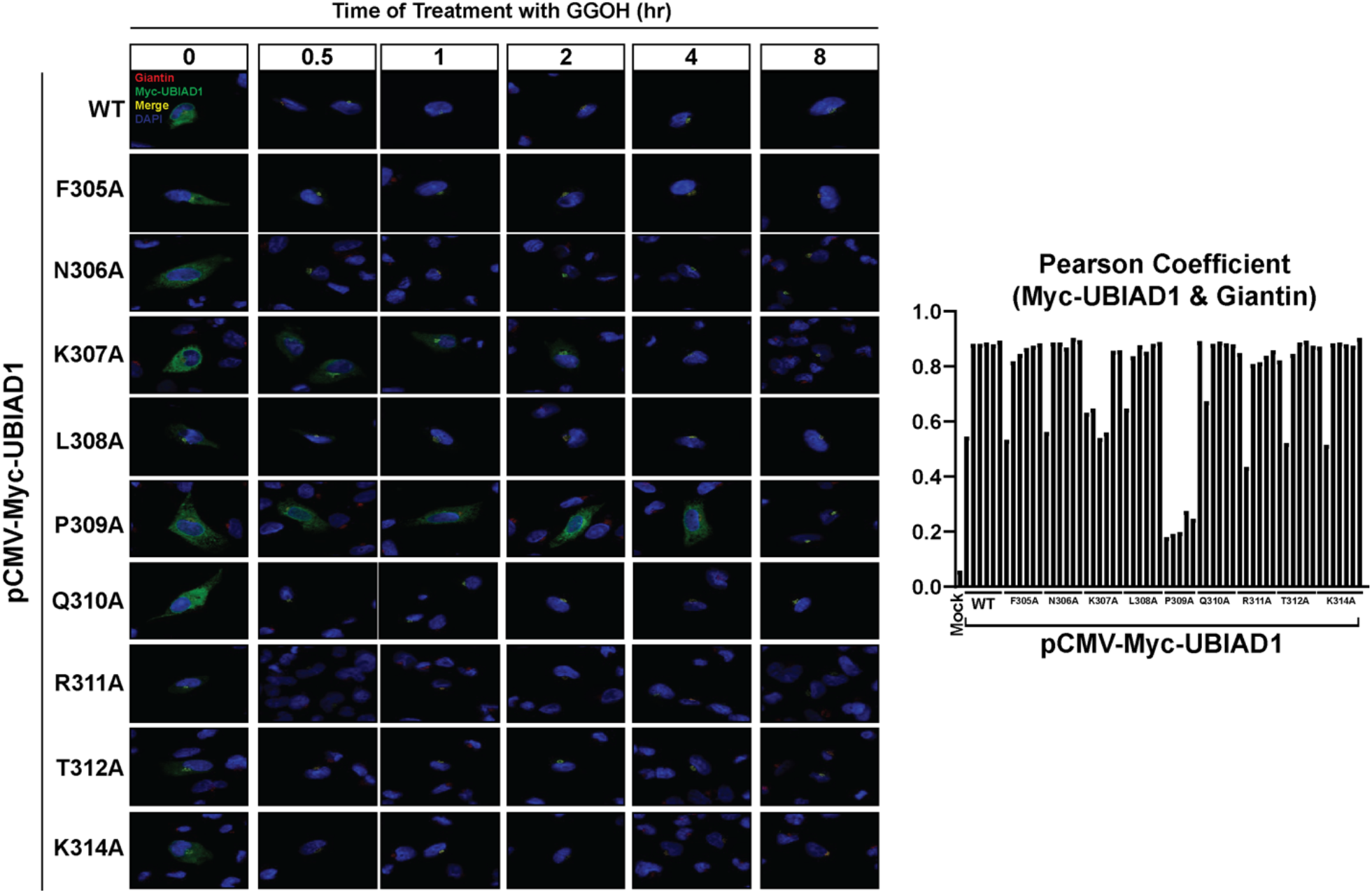
Geranylgeranyl-induced transport of UBIAD1 variants harboring alanine substitutions in cytosolic loop between transmembrane domains 8 and 9. SV-589 cells were set up for experiments on day 0 at 60,000 cells per well in 6 well dish with 2x 12mm coverslip in 5% FCS. Cells were transfected with 0.3 μg DNA per dish using X-tremeGENE HP and sterol depleted overnight. Cells were sterol depleted in media containing 10% NCLPPS, 10 μM compactin and 50 μM mevalonate. Cells were then incubated in media with depleted media and 20 μM GGOH at 30 min, 1hour, 2 hour, 4 hour, and 8 hour time points or left to remain in sterol depleted media. Following incubations described in the figure legends, cells were washed with PBS and subsequently fixed and permeabilized for 15 min in methanol at −20°C. Upon blocking with 1 mg/ml BSA in PBS, coverslips were incubated for 1 hour at room temperature with primary antibodies (rabbit polyclonal anti-GM130 IgG and IgG-9E10, a mouse monoclonal antibody against c-Myc purified from the culture medium of hybridoma clone 9E10 [American Type Culture Collection, Manassas, VA]) diluted in PBS containing 1 mg/ml BSA. Bound antibodies were visualized with goat anti-mouse IgG conjugated to Alexa Fluor 488 and goat anti-rabbit Alexa Fluor 594 (Life Technologies, Grand Island, NY) as described in the figure legends. In addition, coverslips were stained for 5 min with 1 μg/ml 300 nM 4′,6-diamidino-2-phenylindole (Life Technologies) to visualize nuclei. The coverslips were then mounted in Mowiol 4-88 solution (Calbiochem/EMD Millipore, Billerica, MA) and fluorescence analysis was performed using a Plan-Apochromat 63×/1.4 objective (Zeiss, Peabody, MA), an Axio Observer microscope (Zeiss), an Axiocam (Zeiss) color digital camera in black and white mode, and ImageJ software (National Institutes of Health, Bethesda, MD). Brightness levels were adjusted across entire images using ImageJ. The values are the average of 6-10 images (± standard error). The p values were calculated using the Student’s t test: ***, p ≤0.001.

## Discussion

The current data supports the model shown in **Figure 1** for the membrane orientation of UBIAD1, a prenyltransferase that synthesizes vitamin K_2_. For this enzyme, topographical mapping was done using two different methodologies: protease protection and cysteine derivitization. The protease protection experiments demonstrate that the N-terminus is sensitive to protease activity while the C-terminus is not. This indicates that the N-terminus faces the cytosol while the C-terminus faces the lumen. The hydrophobicity plot suggests that UBIAD1 has seven or nine transmembrane domains. Using cysteine derivatization, we determined that UBIAD1 has nine transmembrane domains. This result is consistent with previous studies that determined the cryogenic electron microscopy of SCD-associated UBIAD1 in complex with HMGCR^44^.

When analyzing the subcellular localization of UBIAD1 variants used for cysteine derivatization studies, we noticed that the variant harboring a cysteine substation for G177, L308, and P309 remained sequestered in the ER of isoprenoid-replete cells (**Figure 4**). ER sequestration of the G177C variant can be explained by the observation that G177 is known to be mutated in SCD^45^ and lies near the active site of UBIAD1. To our knowledge, mutation of L308 or P309 have not been observed in SCD families; both residues localize to the cytosolic loop between transmembrane domains 8 and 9 (L8-9), which do not participate in enzymatic activity. This finding prompted us to introduce alanine substitutions of all amino acids in L8-9 of WT UBIAD1 and assess their subcellular localization in response to GGpp. The time course study in **Figure 8** shows that GGpp stimulated Golgi transport of all alanine substitution mutants except P309A. Considering that mutation of P309 renders UBIAD1 refractory to GGpp-induced transport to the Golgi and the residue is not localized to the enzyme’s active site, we conclude that it may be involved in binding to COPII^32^.

A longstanding project in our group has been determining the mechanism through which GGpp induces the translocation of UBIAD1 from the ER to the Golgi. ^2^,^9^,^33^Our current model asserts that UBIAD1 continuously cycles between ER and the Golgi to monitor levels of GGpp embedded in the ER. The binding of GGpp to UBIAD1 induces a conformational change that exposes a binding site for Sec24 and promotes its association with COPII. It is also possible that GGpp binding to UBIAD1 allows binding to an escort protein that in turn facilitates the incorporation of UBIAD1 into COPII vesicles. Cholesterol mediated ER retention of Scap, a protein that binds to sterol regulatory element binding proteins (SREBPs) and mediates their transport from the ER to the Golgi, is another example of a product from the mevalonate pathway controlling the export of a protein from the ER. ^34^,^35^,^36^,^37^ These studies identified a hexapeptide sequence, MELADL, in the cytosolic loop between transmembrane domains six and seven of Scap. This MELADL sequence binds to Sec24 and is necessary for Scap to be incorporated into COPII coated vesicles. When Scap binds to cholesterol, a conformational change occurs that prevents the MELADL sequence from being able to bind to Sec24. This prevents Scap from binding to Sec24 and being incorporated into CopII coated vehicles and instead sequesters Scap to the ER.^38^,^39^ Similarly, we believe that the binding of GGpp to the membrane embedded active site of UBIAD1 induces a conformational change that exposes a binding site for Sec24 and promotes its association with COPII. Our current results indicate that P309 plays a role in this association, but whether this role is direct or indirect remains to be determined. Another possibility is that the putative GGpp-induced conformational change in UBIAD1 allows binding to an escort protein that mediates its incorporation into COPII coated vesicles. We suspect that this is not the case because when UBIAD1 is over expressed it maintains its metabolically regulated transport between the ER and the Golgi. Studies are now underway to determine whether UBIAD1 binds to the Sec23/24 complex and how P309 influences the association. Elucidating the mechanism through which GGpp induces the translocation of UBIAD1 from the ER and Golgi will provide key insight into mechanism governing HMGCR ERAD and utimatley, cholesterol synthesis.

## Methods and Materials

We obtained Map-Peg (QBD10406) from Millipore-Sigma (St. Louis, MO), GGOH (sc-200858) from Santa Cruz Biotechnology (Dallas, TX); MK-4 (V9378) and menadione (M5625) from Sigma-Aldrich (St. Louis, MO). GGpp (I-0200) was purchased from Echelon Bioscience (Salt Lake City, UT), [3H]menadione and [14C]cholesterol were obtained from American Radiolabeled Chemicals (St. Louis, MO). Other reagents including FCS, sodium, compactin, and sodium mevalonate were prepared or obtained as previously described.

### Expression plasmids

The expression plasmid pCMV-Myc-UBIAD1, which encodes human UBIAD1 containing a single copy of a Myc epitope at the N-terminus under transcriptional control of the cytomegalovirus (CMV) promoter, was previously described. The remaining topology mutants of UBIAD1 were generated using the QuikChange® site-directed mutagenesis kit (Agilent Technologies, Santa Clara, CA) and pCMV-Myc-UBIAD1 as a template.

### Cell culture

SV-589 cells are a line of immortalized human fibroblasts expressing the SV40 large T-antigen. Monolayers of SV-589 cells were maintained in medium A (DMEM containing 1,000 mg/l glucose, 100 U/ml penicillin, and 100 μg/ml streptomycin sulfate) supplemented with 10% (v/v) FCS at 37°C, 5% CO2. SV-589(ΔUBIAD1) cells, a line of UBIAD1-deficient SV-589 cells, were grown in medium A supplemented with 10% FCS and 1 mm mevalonate. Chinese hamster ovary (CHO)-K1 cells were grown in medium B (1:1 mixture of Ham’s F-12 medium and Dulbecco’s modified Eagle’s medium containing 100 units/ml penicillin and 100 μg/ml streptomycin sulfate) that contained 5% (v/v) FCS. Monolayers of CHO-K1 cells were grown at 37°C in an 8% CO2 incubator.

### Transfection

SV589 cells were transfected with the X-treme Gene reagent. On day 0, SV589 cells set up at a density of 250,000 cells in 60 mm dishes. On day 1, cells were transfected with 3 μg DNA/dish using a 3:1 ratio of transfection reagent to DNA. X-treme Gene was diluted in media without antibiotics and incubated for 5 min at room temperature prior to being added to the DNA solution. This mixture was then further incubated for 30 min at room temperature. Plates were washed one time with 2 ml of media supplemented with 5% fetal calf serum and refed with 3 ml of the same medium. The X-treme Gene/DNA mixture (0.3 ml) was then added to each dish. Cells were incubated at 37 °C for 16–24 h. On Day 2, the cells were refeed with fresh media supplemented with 5% fetal calf serum. On day 3, the cells were harvested. At the end of the incubation, triplicate dishes of cells for each variable were harvested and pooled for analysis.

CHO-K1 cells were transfected with the Fugene 6 reagent. On day 0, CHO-K1 cells were set up at a density of 250,000 cells in 100 mm dishes. On day 2, cells were transfected with 5μg DNA/dish using a 3:1 ratio of transfection reagent to DNA. Fugene 6 was diluted in media with antibiotics and incubated for 5 min at room temperature prior to being added to the DNA solution. This mixture was then further incubated for 30 min at room temperature. Plates were washed one time with 3 ml of media supplemented with 10% fetal calf serum and refed with 5 ml of the same medium. The Fugene/DNA mixture (1 ml) was then added to each dish. Cells were incubated at 37 °C for 16–24 h. On Day 3, the cells were harvested. At the end of the incubation, triplicate dishes of cells for each variable were harvested and pooled for analysis.

### Subcellular fractionation and immunoblot analysis

Cells were set up for experiments on day 0 as described in the figure legends. After incubations, triplicate dishes were harvested and pooled for analysis. Subcellular fractionation of cells by differential centrifugation was performed as previously described, and aliquots of the resulting membrane fractions were subjected to SDS-PAGE and immunoblot analysis. After SDS-PAGE, proteins were transferred to Hybond C-Extra nitrocellulose filters (GE Healthcare). Primary antibodies used for immunoblot analysis included: rabbit polyclonal anti-calnexin IgG (Novus Biologicals, Littleton, CO), IgG-9E10, a mouse monoclonal antibody against c-Myc purified from the culture medium of hybridoma clone 9E10 (American Type Culture Collection) Mouse monoclonal anti-T7 tag IgG was obtained from EMD Biosciences (San Diego, CA). Bound antibodies were visualized with peroxidase-conjugated, affinity-purified donkey anti-mouse or anti-rabbit IgG (Jackson ImmunoResearch Laboratories, Inc., West Grove, PA) using the SuperSignal CL-HRP substrate system (ThermoFisher Scientific) according to the manufacturer’s instructions. Gels were calibrated with prestained molecular mass markers (Bio-Rad). Filters were exposed to film at room temperature and exposed in a Licor-Fc at room temperature.

### Protease Protection Assay

SV589 cells were transfected with 5 μg of pCMV-MYC-UBIAD1 or pCMV-UBIAD1-FLAG in 100 mm dishes. Microsomes were harvested without protease inhibitors. An equal concentration of microsomal protein was suspended in a total volume of 56 μl in Buffer A and was incubated with the indicated amounts of trypsin for 30 min at 30 °C. Reactions were terminated by the addition of loading buffer and heat inactivation at 95 °C for 10 min.

### PEGylation Assays

The PEGylation method was adapted from Lowe et al. Microsomal protein (20 μg) was suspended in a total volume of 54 μl of Buffer A containing protease inhibitors. Microsomes were treated with or without 1% (v/v) Triton X-100 or SDS as indicated. Microsomes were then treated with or without mPEG (catalog number 10406, Quanta Biodesign) (5 mm, 30 min) at 37 °C. Reactions were quenched with dithiothreitol (DTT) (10 mm, 10 min, room temperature). Samples were subjected to Novex 4-20% Tris-Glycine followed by Western blotting.

### Synthesis of [3H]MK-4 in intact cells

To measure [3H]MK-4 synthesis in intact cells, we set up cells on day 0 as described in the figure legends. On day 2, the cells were refed medium A with 5% FCS and 2.5 μCi/ml [3H]menadione (unlabeled menadione was added to achieve a final concentration of 50 nM). Following incubation for 16 h at 37°C, cells were washed twice with PBS containing 2% BSA, followed by an additional wash with PBS. The cells were lysed with 0.1 N NaOH; an aliquot of each lysate was removed for protein determination. The remaining lysates were mixed with recovery solution containing 16 μg/ml MK-4, 0.025 μCi/ml [14C]cholesterol, and 16 μg/ml unlabeled cholesterol and extracted with dichloromethane:methanol (2:1). The resulting lipids were subjected to TLC and incorporation of [3H]menadione into MK-4 was determined as previously described.

### Immunofluorescence

SV-589 cells were set up for experiments on day 0 as described in the figure legends. Following incubations described in the figure legends, cells were washed with PBS and subsequently fixed and permeabilized for 15 min in methanol at −20°C. Upon blocking with 1 mg/ml BSA in PBS, coverslips were incubated for 1 hour at room temperature with primary antibodies (rabbit polyclonal anti-GM130 IgG and IgG-9E10, a mouse monoclonal antibody against c-Myc purified from the culture medium of hybridoma clone 9E10 [American Type Culture Collection, Manassas, VA]) diluted in PBS containing 1 mg/ml BSA. Bound antibodies were visualized with goat anti-mouse IgG conjugated to Alexa Fluor 488 and goat anti-rabbit Alexa Fluor 594 (Life Technologies, Grand Island, NY) as described in the figure legends. In addition, coverslips were stained for 5 min with 1 μg/ml 300 nM 4′,6-diamidino-2-phenylindole (Life Technologies) to visualize nuclei. The coverslips were then mounted in Mowiol 4-88 solution (Calbiochem/EMD Millipore, Billerica, MA) and fluorescence analysis was performed using a Plan-Apochromat 63×/1.4 objective (Zeiss, Peabody, MA), an Axio Observer microscope (Zeiss), an Axiocam (Zeiss) color digital camera in black and white mode, and ImageJ software (National Institutes of Health, Bethesda, MD).

### Stability Assay

SV-589 cells were set up for experiments on day 1 400K cells per 100-mm dish in media containing 5% FCS. On day 1, cells were transfected with 3 μg/dish of pCMV-HMG-Red(TM1-8)-T7 together with 100 ng of pCMV-Myc-UBIAD1 in media containing 5% FCS. Following incubation for 16 hr at 37°C, cells were harvested and subjected to subcellular fractionation. Aliquots of resulting membrane fractions were then subjected to SDS-PAGE and immunoblot analysis was carried out with anti-T7 IgG (against reductase), IgG-9E10 (against UBIAD1), and anti-calnexin IgG.

## Acknowledgements

We thank Lisa Beatty for help with tissue culture. This work was supported by the NIH grants R01 GM134700, GM144039, and P01 HL160487 (to R.D.B).

## Supplemental Data

**Supplemental Figure 1:**
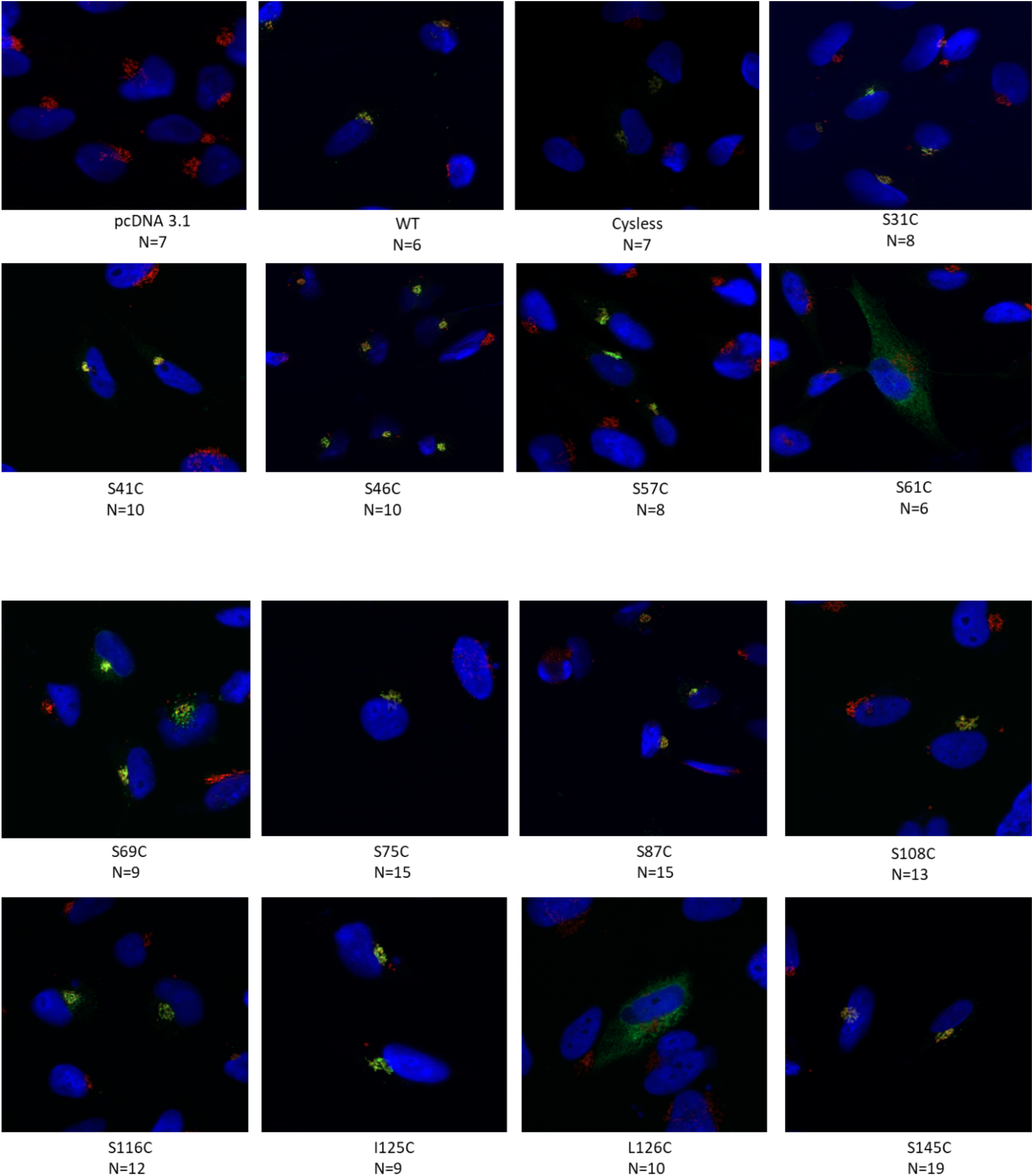
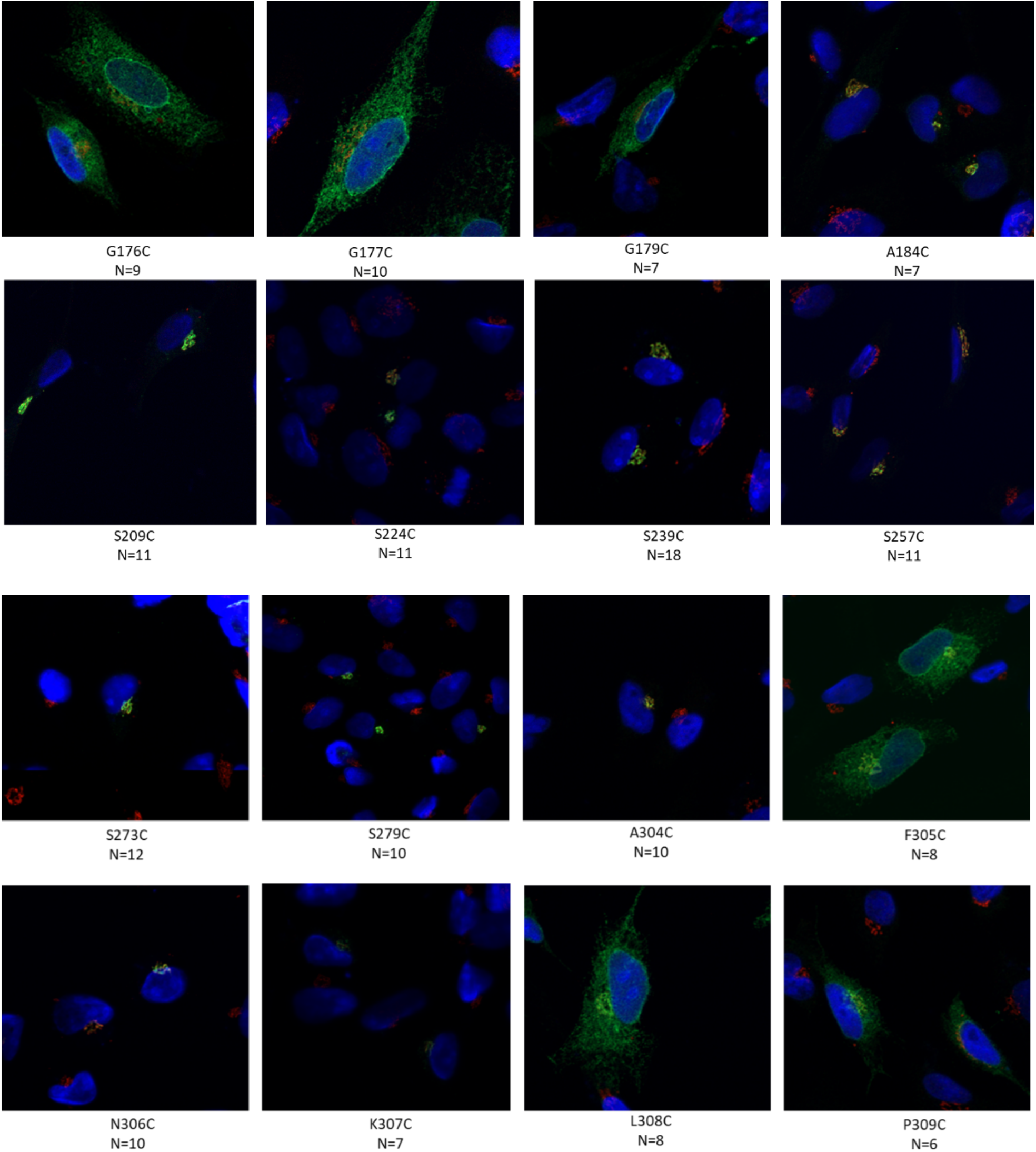
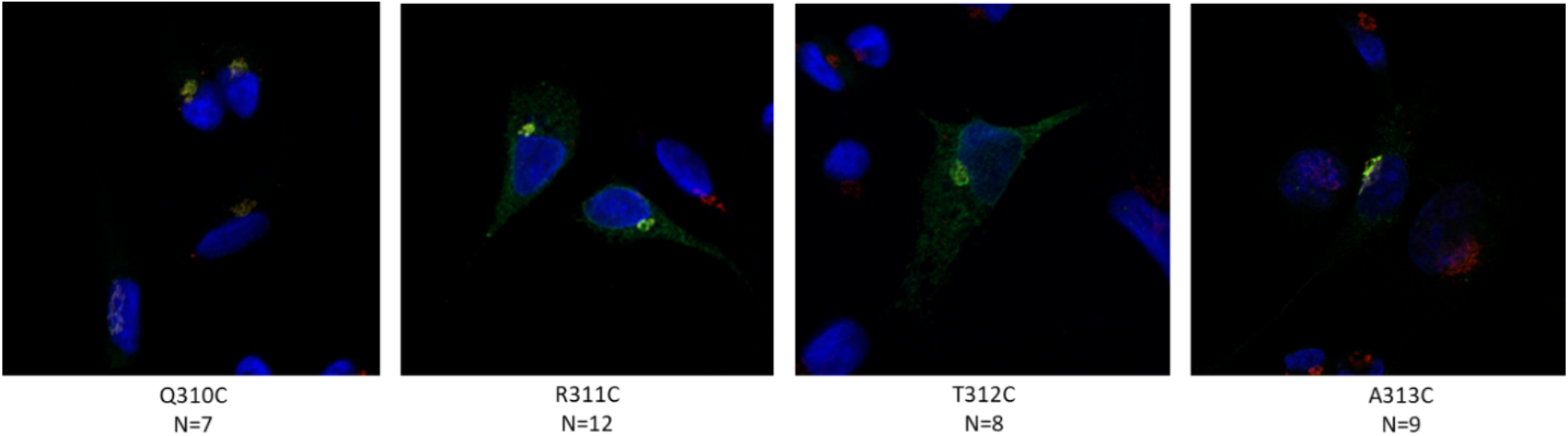
Immunofluorescent Images of topology mutants with number of cells analyzed for Pearson Coefficient measurements. SV-589 cells were set up for experiments on day 0 at 150K in 60 mm dishes in 5% FCS. On day 1, the cells were transfected with 0.5 μg/ml pf plasmid DNA in 5% FCS. On Day 3, the cells were washed with PBS and subsequently fixed and permeabilized for 15 min in methanol at −20°C. Upon blocking with 1 mg/ml BSA in PBS, coverslips were incubated for 1 hour at room temperature with primary antibodies (rabbit polyclonal anti-GM130 IgG and IgG-9E10, a mouse monoclonal antibody against c-Myc purified from the culture medium of hybridoma clone 9E10 [American Type Culture Collection, Manassas, VA]) diluted in PBS containing 1 mg/ml BSA. Bound antibodies were visualized with goat anti-mouse IgG conjugated to Alexa Fluor 488 and goat anti-rabbit Alexa Fluor 594 (Life Technologies, Grand Island, NY) as described in the figure legends. In addition, coverslips were stained for 5 min with 1 μg/ml 300 nM 4′,6-diamidino-2-phenylindole (Life Technologies) to visualize nuclei. The coverslips were then mounted in Mowiol 4-88 solution (Calbiochem/EMD Millipore, Billerica, MA) and fluorescence analysis was performed using a Plan-Apochromat 63×/1.4 objective (Zeiss, Peabody, MA), an Axio Observer microscope (Zeiss), an Axiocam (Zeiss) color digital camera in black and white mode, and ImageJ software (National Institutes of Health, Bethesda, MD). Brightness levels were adjusted across entire images using ImageJ. The values are the average of 6-10 images (± standard error). The p values were calculated using the Student’s t test: ***, p ≤0.001.

**Supplemental Figure 2:**
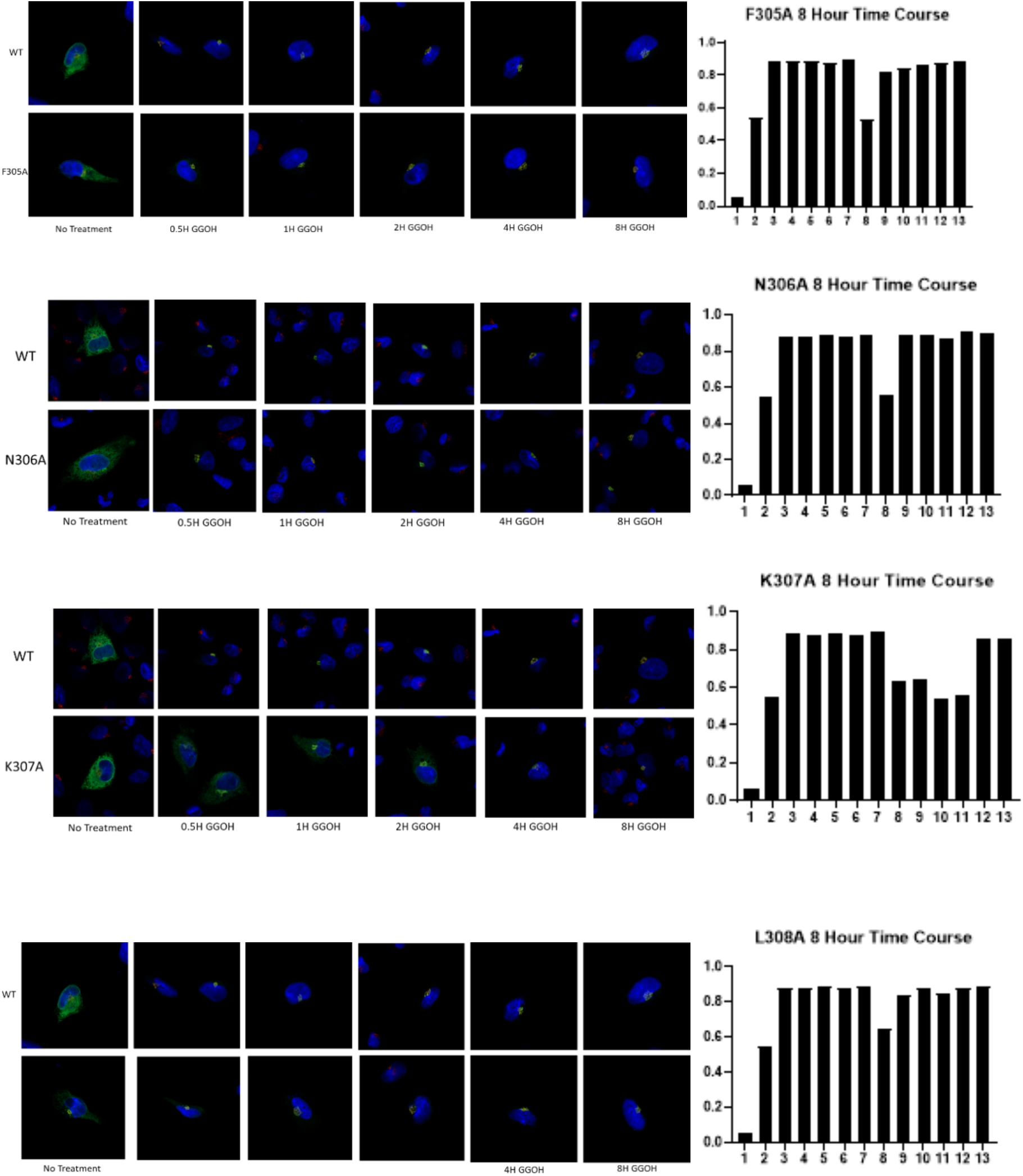
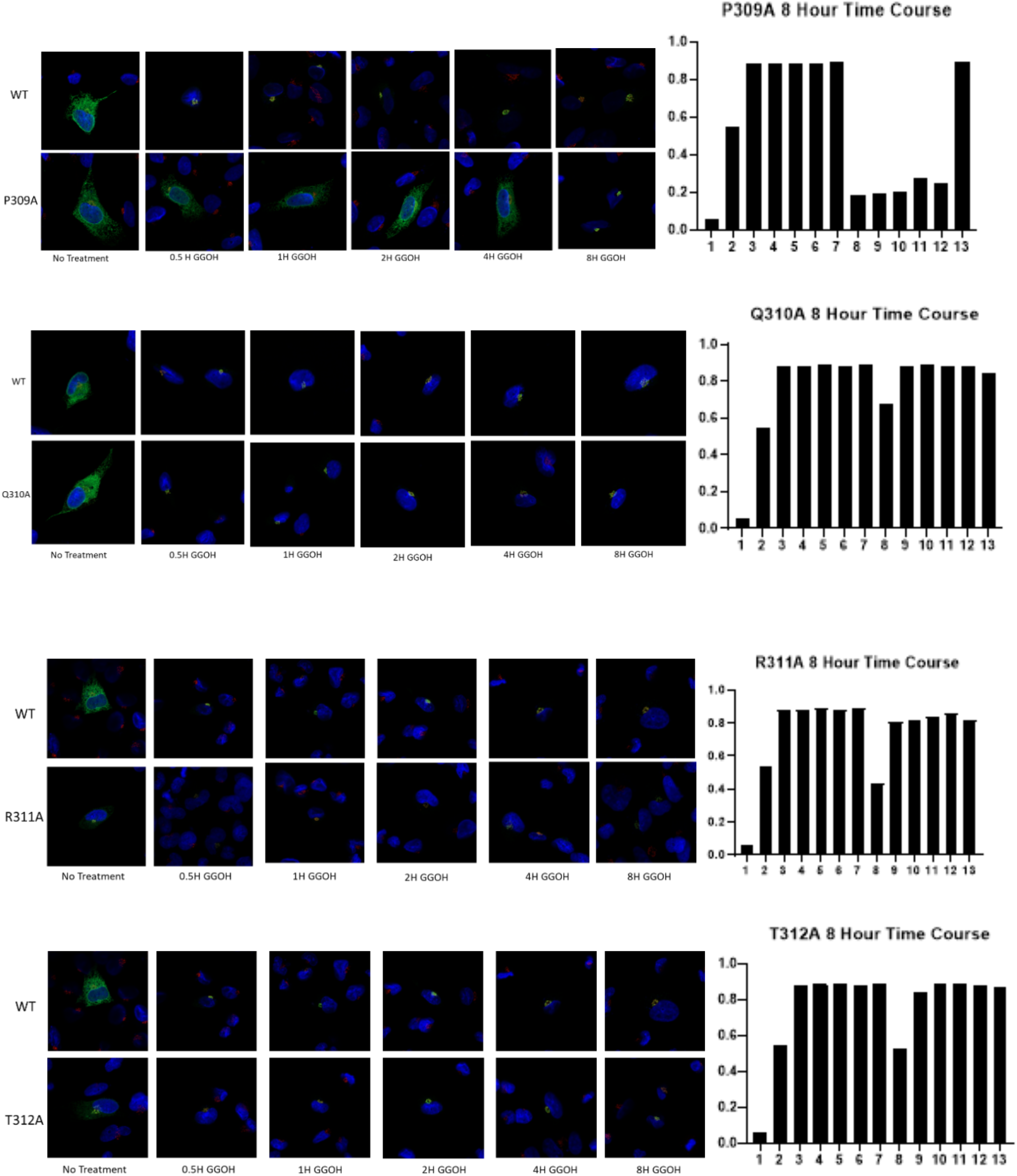
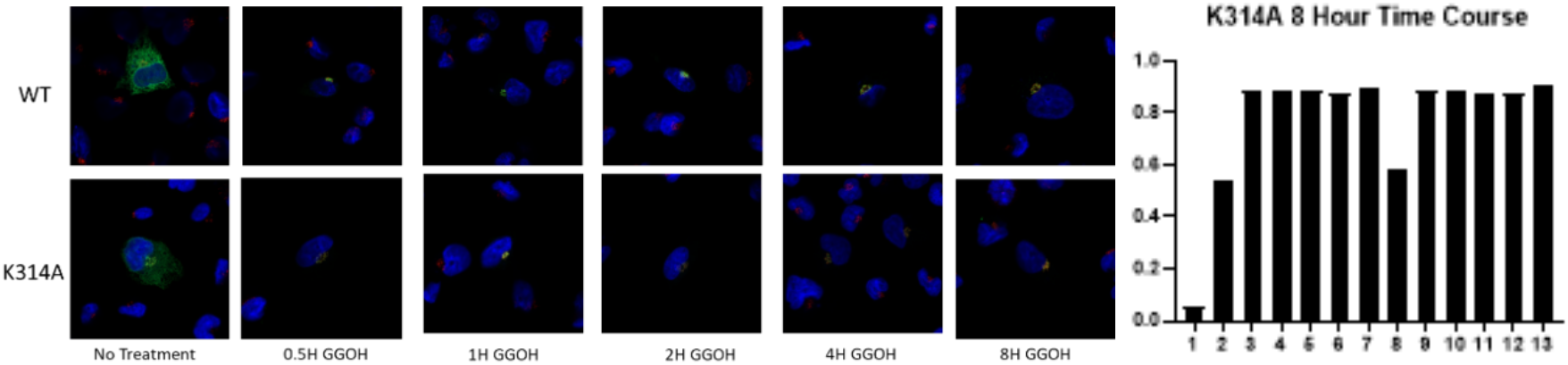
Individual Experiments of Alanine Scan Eight Hour Time Course demonstrating geranylgeranyl-induced transport of UBIAD1 variants harboring alanine substitutions in cytosolic loop between transmembrane domains 8 and 9. SV-589 cells were set up for experiments on day 0 at 60,000 cells per well in 6 well dish with 2x 12mm coverslip in 5% FCS. Cells were transfected with 0.3 μg DNA per dish using X-tremeGENE HP and sterol depleted overnight. Cells were sterol depleted in media containing 10% NCLPPS, 10 μM compactin and 50 μM mevalonate. Cells were then incubated in media with depleted media and 20 μM GGOH at 30 min, 1hour, 2 hour, 4 hour, and 8 hour time points or left to remain in sterol depleted media. Following incubations described in the figure legends, cells were washed with PBS and subsequently fixed and permeabilized for 15 min in methanol at −20°C. Upon blocking with 1 mg/ml BSA in PBS, coverslips were incubated for 1 hour at room temperature with primary antibodies (rabbit polyclonal anti-GM130 IgG and IgG-9E10, a mouse monoclonal antibody against c-Myc purified from the culture medium of hybridoma clone 9E10 [American Type Culture Collection, Manassas, VA]) diluted in PBS containing 1 mg/ml BSA. Bound antibodies were visualized with goat anti-mouse IgG conjugated to Alexa Fluor 488 and goat anti-rabbit Alexa Fluor 594 (Life Technologies, Grand Island, NY) as described in the figure legends. In addition, coverslips were stained for 5 min with 1 μg/ml 300 nM 4′,6-diamidino-2-phenylindole (Life Technologies) to visualize nuclei. The coverslips were then mounted in Mowiol 4-88 solution (Calbiochem/EMD Millipore, Billerica, MA) and fluorescence analysis was performed using a Plan-Apochromat 63×/1.4 objective (Zeiss, Peabody, MA), an Axio Observer microscope (Zeiss), an Axiocam (Zeiss) color digital camera in black and white mode, and ImageJ software (National Institutes of Health, Bethesda, MD). Brightness levels were adjusted across entire images using ImageJ. The values are the average of 6-10 images (± standard error). The p values were calculated using the Student’s t test: ***, p ≤ 0.001.

**Supplemental Figure 3:**
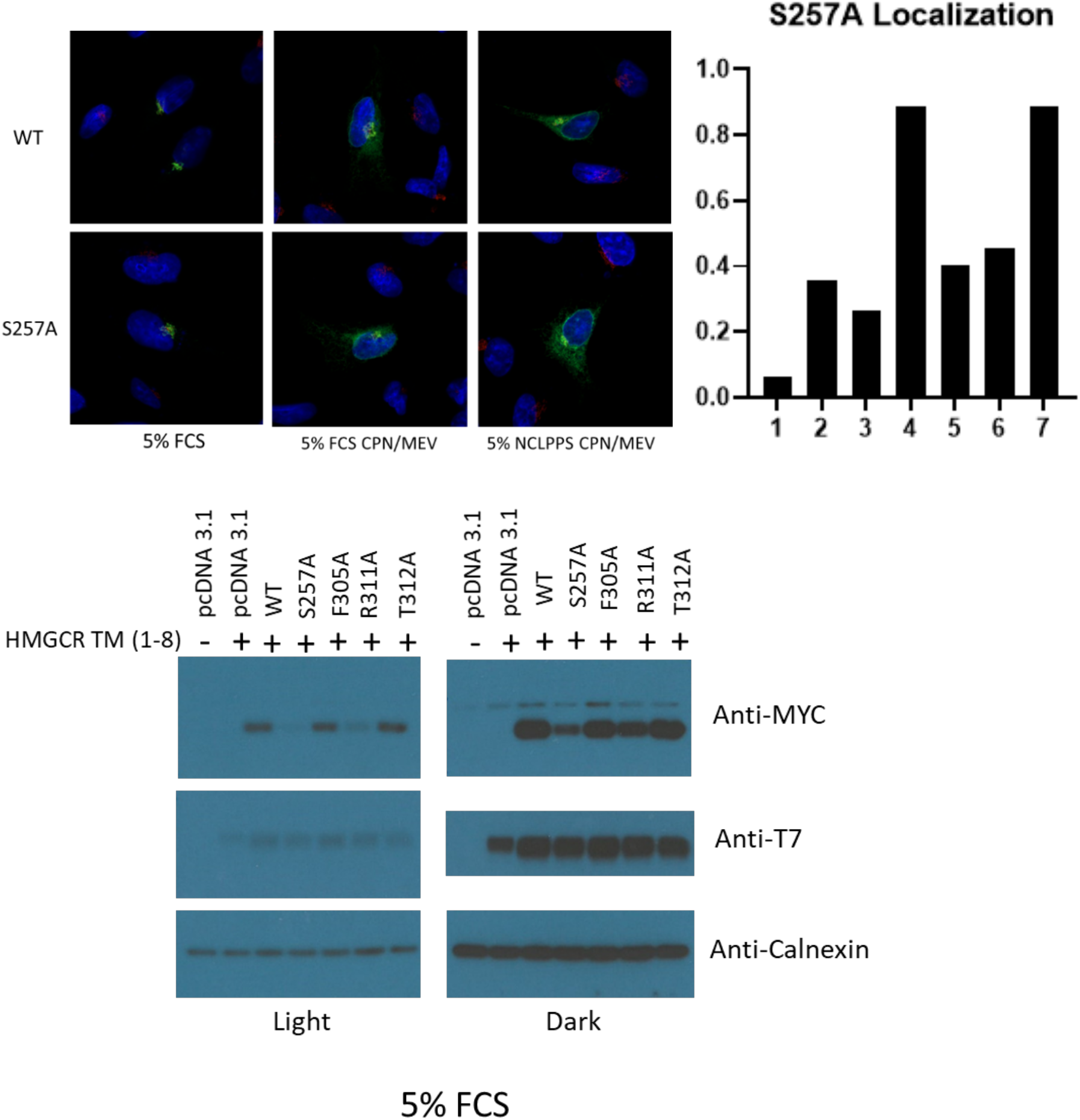
Follow up experiments on topology mutants that do not stabilize HMGCR or hyperstabilize HMGCR in a WT background. **Subcellular localization (A) and HMG CoA reductase stabilization (B) of UBIAD1 variants harboring alanine substitutions in cytosolic loop between transmembrane domains 8 and 9 that hyperstabilize or fail to stabilize HMGCR in the cysteineless background. A**, SV-589 cells were set up for experiments on day 0 at 150K in 60 mm dishes in 5% FCS. On day 1, the cells were transfected with 0.5 μg/ml pf plasmid DNA in 5% FCS. On Day 3, the cells were washed with PBS and subsequently fixed and permeabilized for 15 min in methanol at −20°C. Upon blocking with 1 mg/ml BSA in PBS, coverslips were incubated for 1 hour at room temperature with primary antibodies (rabbit polyclonal anti-GM130 IgG and IgG-9E10, a mouse monoclonal antibody against c-Myc purified from the culture medium of hybridoma clone 9E10 [American Type Culture Collection, Manassas, VA]) diluted in PBS containing 1 mg/ml BSA. Bound antibodies were visualized with goat anti-mouse IgG conjugated to Alexa Fluor 488 and goat anti-rabbit Alexa Fluor 594 (Life Technologies, Grand Island, NY) as described in the figure legends. In addition, coverslips were stained for 5 min with 1 μg/ml 300 nM 4′,6-diamidino-2-phenylindole (Life Technologies) to visualize nuclei. The coverslips were then mounted in Mowiol 4-88 solution (Calbiochem/EMD Millipore, Billerica, MA) and fluorescence analysis was performed using a Plan-Apochromat 63×/1.4 objective (Zeiss, Peabody, MA), an Axio Observer microscope (Zeiss), an Axiocam (Zeiss) color digital camera in black and white mode, and ImageJ software (National Institutes of Health, Bethesda, MD). Brightness levels were adjusted across entire images using ImageJ. The values are the average of 6-10 images (± standard error). **B**, SV-589 cells were set up for experiments on day 1 400K cells per 100-mm dish in containing 5% FCS. On day 1, cells were transfected with 3 μg/dish of pCMV-HMG-Red(TM1-8)-T7 together with 100 ng of pCMV-Myc-UBIAD1 in media containing 5% FCS. Following incubation for 16 hr at 37°C, cells were harvested and subjected to subcellular fractionation. Aliquots of resulting membrane fractions were then subjected to SDS-PAGE and immunoblot analysis was carried out with anti-T7 IgG (against reductase), IgG-9E10 (against UBIAD1), and anti-calnexin IgG.

